# Evaluating the reliability of human brain white matter tractometry

**DOI:** 10.1101/2021.02.24.432740

**Authors:** John Kruper, Jason D. Yeatman, Adam Richie-Halford, David Bloom, Mareike Grotheer, Sendy Caffarra, Gregory Kiar, Iliana I. Karipidis, Ethan Roy, Bramsh Q. Chandio, Eleftherios Garyfalldis, Ariel Rokem

## Abstract

The validity of research results depends on the reliability of analysis methods. In recent years, there have been concerns about the validity of research that uses diffusion-weighted MRI (dMRI) to understand human brain white matter connections *in vivo*, in part based on reliability of the analysis methods used in this field. We defined and assessed three dimensions of reliability in dMRI-based tractometry, an analysis technique that assesses the physical properties of white matter pathways: (1) reproducibility, (2) test-retest reliability and (3) robustness. To facilitate reproducibility, we provide software that automates tractometry (https://yeatmanlab.github.io/pyAFQ). In measurements from the Human Connectome Project, as well as clinical-grade measurements, we find that tractometry has high test-retest reliability that is comparable to most standardized clinical assessment tools. We find that tractometry is also robust: showing high reliability with different choices of analysis algorithms. Taken together, our results suggest that tractometry is a reliable approach to analysis of white matter connections. The overall approach taken here both demonstrates the specific trustworthiness of tractometry analysis and outlines what researchers can do to demonstrate the reliability of computational analysis pipelines in neuroimaging.

## Introduction

The white matter of the brain contains the long-range connections between distant cortical regions. The integration and coordination of brain activity through the fascicles containing these connections is important for information processing and for brain health (1,2). Using voxel-specific directional diffusion information from diffusion-weighted MRI (dMRI), computational tractography produces three-dimensional trajectories through the white matter within the MRI volume that are called “streamlines”(3,4). Collections of streamlines that match the location and direction of major white matter pathways within an individual can be generated with different strategies: using probabilistic (5,6) or streamline-based (7,8) atlases, or known anatomical landmarks (9–12). Because these are models of the anatomy, we refer to these estimates as “bundles” to distinguish them from the anatomical pathways themselves. The delineation of well-known anatomical pathways overcomes many of the concerns about confounds in dMRI-based tractography (13,14), because “brain connections derived from diffusion MRI tractography can be highly anatomically accurate – if we know where white matter pathways start, where they end, and where they do not go”(15).

The physical properties of the tissue affect the diffusion of water within the brain and the microstructure of tissue within the white matter along the length of computationally-generated bundles can be assessed using a variety of models (16,17). Taken together, computational tractography, bundle recognition and diffusion modeling provide so-called “tract profiles”: estimates of microstructural properties of tissue along the length of major pathways. This is the basis of tractometry: statistical analysis that compares different groups, or assesses individual variability in brain connection structure (9,18–21). For the inferences made from tractometry to be valid and useful, tract profiles need to be reliable.

In the present work, we provide an assessment of three different ways in which scientific results can be reliable: reproducibility, test-retest reliability, and robustness. These terms are often debated and conflicting definitions for these terms have been proposed (22,23). Here, we use the definitions proposed in (24). *Reproducibility* is defined as the case in which data and methods are fully accessible and usable: running the same code with the same data should produce an identical result. Use of different data (e.g., in a test-retest experiment) resulting in quantitatively comparable results would denote *test-retest reliability (TRR)*. In clinical science and psychology in general, TRR (e.g., in the form of inter-rater reliability) is considered a key metric of the reliability of a measurement. Use of a different analysis approach or different analysis system (e.g., different software implementation of the same ideas) could result in similar conclusions, denoting their *robustness* against implementation details. The recent findings of Botvinik-Nezer *et al* (25) show that even when full computational reproducibility is achieved, the results of analysing a single fMRI dataset can vary significantly between teams and analysis pipelines, demonstrating issues of robustness.

The contribution of the present work is three-fold: To support reproducible research using tractometry, we developed an open-source software library called Automated Fiber Quantification in Python (pyAFQ; https://yeatmanlab.github.io/pyAFQ). Given dMRI data that has undergone standard preprocessing (e.g., using QSIprep (26)), pyAFQ automatically performs tractography, classifies streamlines into bundles representing the major tracts, and extracts tract profiles of diffusion properties along those bundles, producing “tidy” CSV output files (27) that are amenable to further statistical analysis (Fig. S1). The library implements the major functionality provided by a previous MATLAB implementation of tractometry analysis (9), and offers a menu of configurable algorithms allowing researchers to tune the pipeline to their specific scientific questions (Fig. S2). Second, we use pyAFQ to assess test-retest reliability of tractometry results. Third, we assess robustness of tractometry results to variations across different models of the diffusion in individual voxels, across different bundle recognition approaches, and across different implementations.

## Materials and Methods

### pyAFQ

We developed an open-source tractometry software library to support computational reproducibility: Python Automated Fiber Quantification (pyAFQ; https://github.com/yeatmanlab/pyAFQ). The software relies heavily on methods implemented in DIPY (28). Our implementation was also guided by a previous MATLAB implementation of tractometry (mAFQ) (9). More details are available in the ‘Automated Fiber Quantification in Python (pyAFQ)’ section of Supplementary Methods.

### Tractometry

The pyAFQ software is configurable, allowing users to specify methods and parameters for different stages of the analysis (Fig. S2). Here, we will describe the default setting. In the first step, computational tractography methods, implemented in DIPY (28), are used to generate streamlines throughout the brain white matter (Fig. S1**A**). Next, the T1-weighted MNI template (29,30) is registered to the anistropic power map (APM) (31,32) computed from the diffusion data, that has a T1-like contrast (Fig. S1**B**) using the symmetric image normalization method (33) implemented in DIPY (28). The next step is to perform bundle recognition, where each tractography streamline is classified as either belonging to a particular bundle, or discarded. We use the transform found during registration to bring canonical anatomical landmarks, such as waypoint regions of interest (ROIs) and probability maps, from template space to the individual subject’s native space. Waypoint ROIs are used to delineate the trajectory of the bundles (34). See Table S1 for the bundle abbreviations we use in this paper. Streamlines that pass through inclusion waypoint ROIs for a particular bundle, and do not pass through exclusion ROI, are selected as candidates to include in the bundle. In addition, a probabilistic atlas (35) is used as a tie-breaker to determine whether a streamline is more likely to belong to one bundle or another (in cases where the streamline matches the criteria for inclusion in either). For example, the corticospinal tract is identified by finding streamlines that do pass through an axial waypoint ROI in the brainstem and another ROI axially oriented in the white matter of the corona radiata, but that do not pass through the midline (Fig. S1**C**). The final step is to extract the tract profile: each streamline is resampled to a fixed number of points and the mean value of a diffusion-derived scalar (e.g., fractional anisotropy (FA) and mean diffusivity (MD)) is found for each one of these nodes. The values are summarized by weighting the contribution of each streamline, based on how concordant the trajectory of this streamline is with respect to the other streamlines in the bundle (Fig. S1**D**). To make sure that profiles represent properties of the core white matter, we remove the first and last 5 nodes of the profile, then further remove any nodes where either the FA is less than 0.2 or the MD is greater than 0.002. This removes nodes that contain partial volume artifacts (16).

### Data

We used two datasets with test-retest measurements. We used Human Connectome Project test-retest measurements of dMRI for 44 neurologically healthy subjects aged 22-35 (HCP-TR) (36). The other is an experimental dataset, with dMRI from 48 children, 5 years old in age, collected at the University of Washington (UW-PREK). More details about the measurement are available in the ‘Data’ section of Supplementary Methods.

### HCP-TR Configurations

We processed HCP-TR with three different pyAFQ configurations. In the first configuration, we used the diffusion kurtosis model (DKI) as the orientation distribution function (ODF) model. In the second configuration, we used constrained spherical deconvolution (CSD) as the ODF model. For the final configuration, we used RecoBundles (8) for bundle recognition instead of the default waypoint ROI approach, and DKI as the ODF model. More details are available in the ‘Configurations’ section of Supplementary Methods.

### Measures of Reliability

Tract recognition of each bundle was compared across measurements and methods using the Dice coefficient, weighted by streamline count (wDSC) (37). Tract profiles were compared with three measures: (1) Profile reliability: mean intraclass correlation coefficient (ICC) across points in different tract profiles for different data, which quantifies the *agreement* of tract profiles (38,39); (2) Subject reliability: Spearman’s rank correlation coefficient (Spearman’s *ρ*) between the mean of the tract profiles across individuals, which quantifies the *consistency* of the mean of tract profiles; (3) an adjusted contrast index profile (ACIP) to directly compare the values of individual nodes in the tract profiles in different measurements. To estimate test-retest reliability (TRR), the above measures were calculated for each individual across different measurements. To estimate robustness, these were calculated for each individual across different analysis methods. For example, if we calculate the subject reliability across analysis methods, we would call that “subject robustness”. If we calculated subject reliability across measurements, we would call that “subject TRR”. We explain profile and subject reliability in more detail below; we explain wDSC and ACIP in more detail in the ‘Measures of Reliability’ section of Supplementary Methods

#### Profile reliability

We use profile reliability to compare the shapes of profiles per bundle and per scalar. Given two sets of data (either test-retest or from different analyses), we first calculate the ICC between tract profiles for each subject in a given bundle and scalar. Then, we take the mean of those correlations. We do this for every bundle and for every scalar. We call this profile reliability because larger differences in the overall values along the profiles will result in a smaller mean of the ICC. Consistent profile shapes are important for distinguishing bundles. Profile reliability provides an assessment of the overall reliability of the tract profiles, summarizing over the full length of the bundle, for a particular scalar. We calculate the 95% confidence interval on profile reliabilities using the standard error of the measurement.

In some cases, there is low between-subject variance in tract profile shape (for example, this is often the case in CST). We use ICC to account for this, as ICC will penalize low between-subject variance in addition to rewarding high within-subject variance. Profile reliability is a way of quantifying the *agreement* between profiles. Qualitatively, we use four descriptions for profile reliability: excellent (ICC > 0.75), good (ICC = 0.60 to 0.74), fair (ICC = 0.40 to 0.59), and poor (ICC < 0.40) (40).

#### Subject reliability

We calculate subject reliability to compare individual differences in profiles, per bundle and per scalar, following (41). Given two measurements for each subject, we first take the mean of each profile within each individual, measurement and scalar. Then we calculate Spearman’s *ρ* from the means from different subjects for a given bundle and scalar across the measurements. High subject reliability means the ordering of an individual’s tract profile mean among other individuals is consistent across measurements or methods. This is akin to test reliability which is computed for any clinical measure.

One downside of subject reliability is that the shape of the extracted profile is not considered. Additionally, if one measurement or method produces higher values for all subjects uniformly, subject reliability would not be affected. Instead, the intent of subject reliability is to well summarize the preservation of relative differences between individuals for mean tract profiles. In other words, subject reliability quantifies the *consistency* of mean profiles. The 95% confidence interval on subject reliabilities are parametric.

## Results

Tractometry using pyAFQ classifies streamlines into bundles that represent major anatomical pathways. The streamlines are used to sample dMRI-derived scalars into bundle profiles that are calculated for every individual and can be summarized for a group of subjects. An example of the process and result of the tract profile extraction process is shown in Supplementary Fig. S3, together with the results of this process across the 18 major white matter pathways for all subjects in the HCP-TR dataset.

### Assessing test-retest reliability of tractometry

In datasets with scan-rescan data we can assess test-retest reliability (TRR) at several different levels of tractometry. For example, the correlation between two profiles provides a measure of the reliability of the overall tract profile in that subject. Analyzing the Human Connectome Project’s test-retest dataset (HCP-TR), we find that for fractional anisotropy (FA) calculated using DKI, the values of *profile reliability* vary across subjects (Figure 1**A**), but they overall tend to be rather high, with the average value within each bundle in the range 0.77 ±0.05 to 0.92 ±0.02 and a median across bundles of 0.86 (Figure 1**B**). We find similar results for mean diffusivity (MD; Fig. S4) and replicate similar results in a second dataset (Fig. 3**B**).

**Fig. 1.**
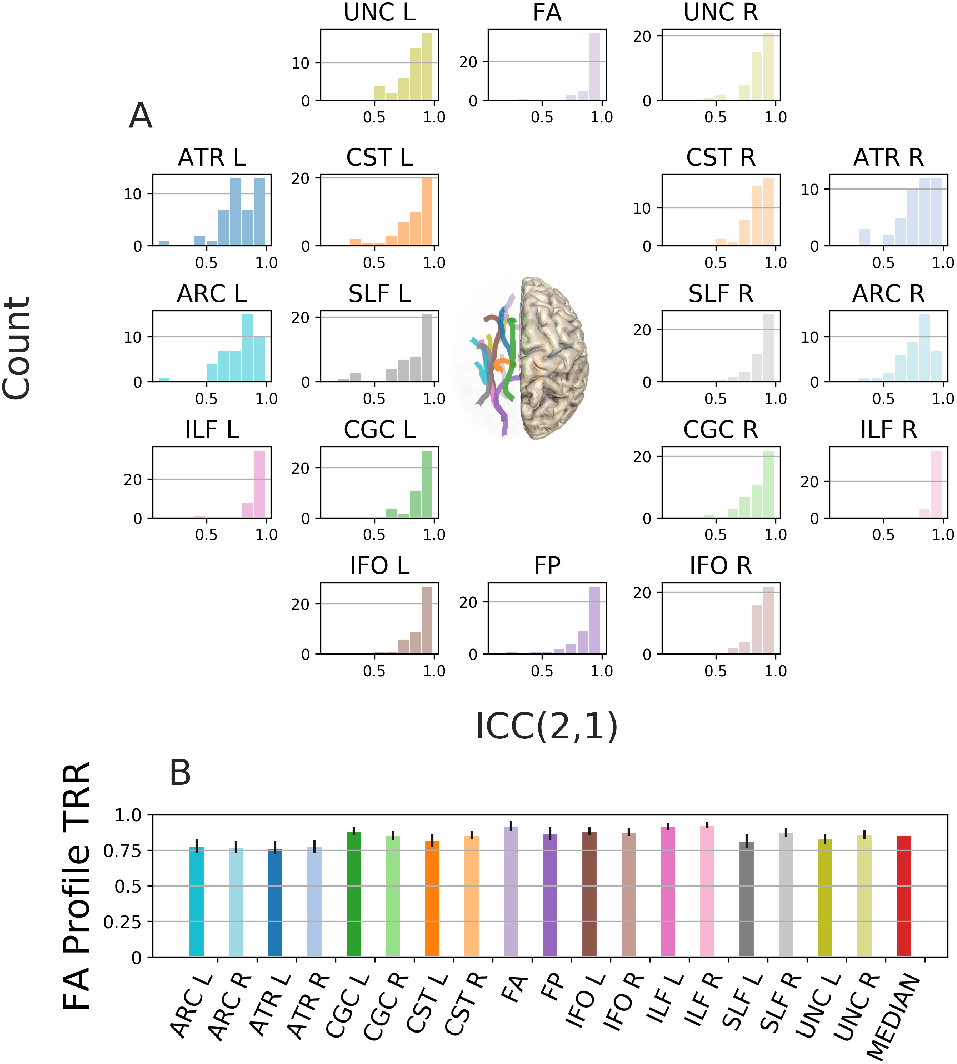
FA profile test-retest reliability. **A:** Histograms of individual subject ICC between the FA tract profiles across sessions for a given bundle. Colors encode the bundles, matching the diagram showing the rough anatomical positions of the bundles for the left side of the brain (center). **B:** Mean (±95% confidence interval) TRR for each bundle, color-coded to match the histograms and the bundles diagram, with median across bundles in red.

**Fig. 2.**
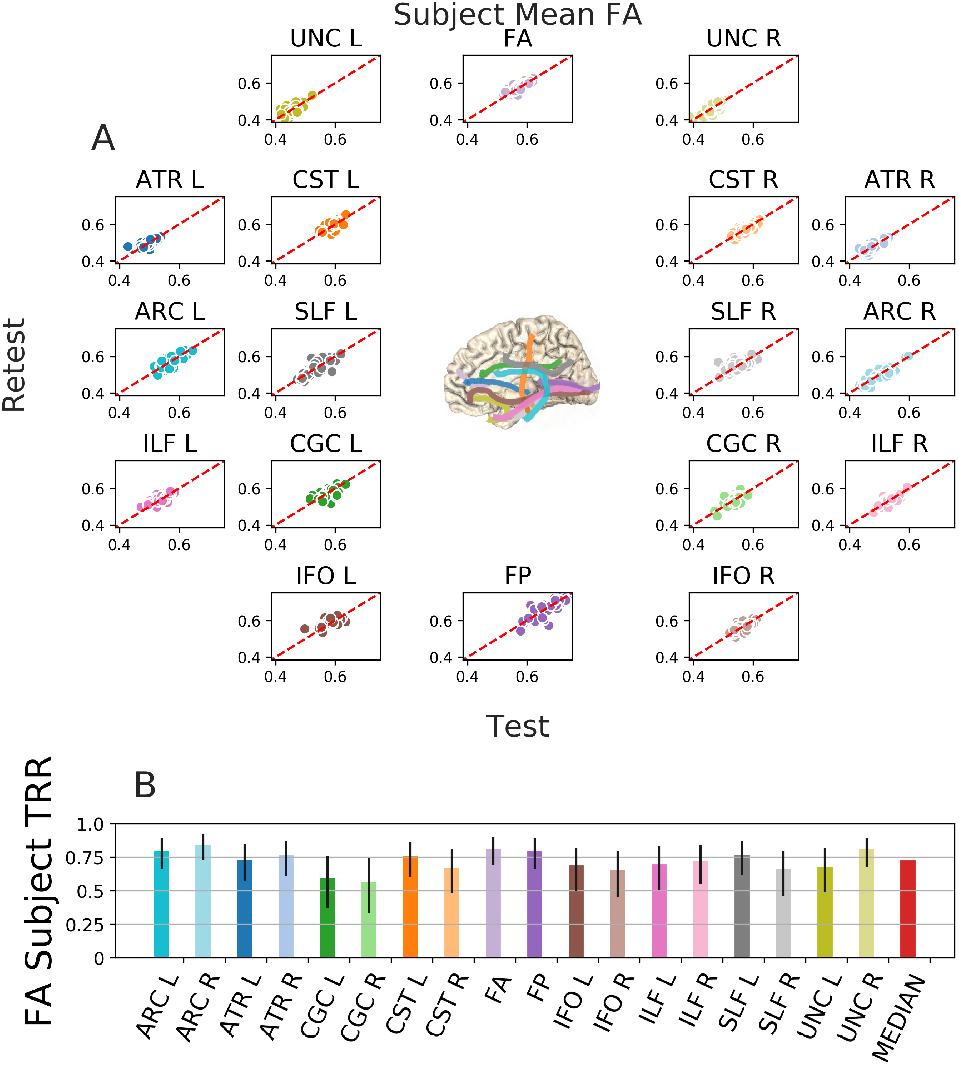
Subject test-retest reliability. **A:** Mean tract profiles for a given bundle and the FA scalar for each subject using the first and second session of HCP-TR. Colors encode bundle information, matching the core of the bundles (center). **B:** subject reliability is calculated from the Spearman’s *ρ* of these distributions, with median across bundles in red (±95% confidence interval).

**Fig. 3.**
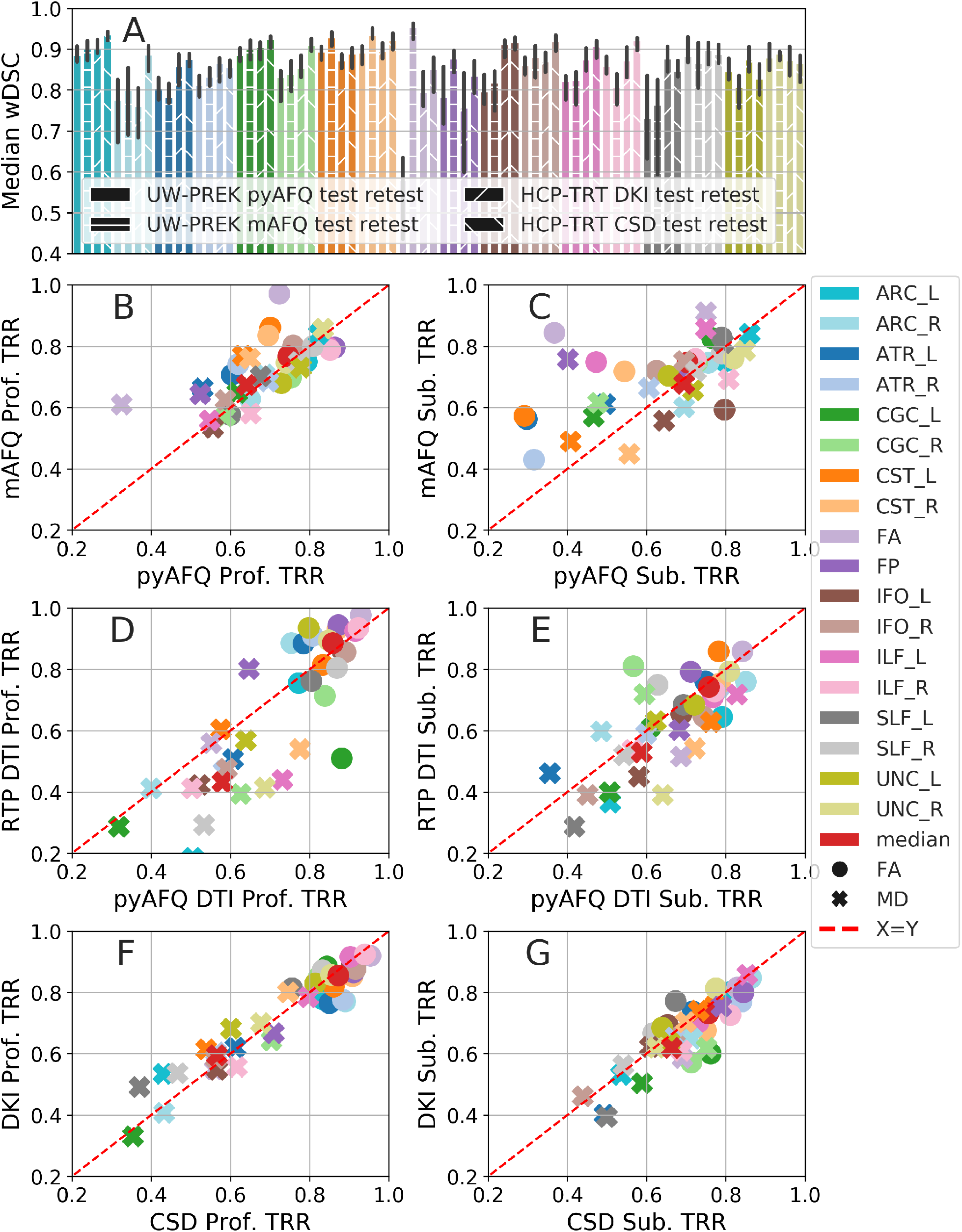
wDSC, profile, and subject TRR of: pyAFQ and mAFQ on UW-PREK; pyAFQ on HCP-TR using different ODF models; and RTP on HCP-TR. Colors indicate bundle. In **A**: texture indicates the dataset and methods being compared. Error bars show the 95% confidence interval. **B**, **D**, and **F** show profile TRR and **C**, **E**, and **G** show subject TRR. Profile and subject TRR calculations are demonstrated with HCP-TR using DKI in figures 1 and 2 respectively. In **B** and **C**, we compare the TRR of mAFQ and pyAFQ on UW-PREK. In **D** and **E**, we compare pyAFQ and RTP on HCP-TR using only single shell data. In **F** and **G**, we compare DKI and CSD TRR on HCP-TR. Point shapes indicate the extracted scalar. The red dotted line is equal TRR between methods.

#### Subject reliability

assesses the reliability of mean tract profiles across individuals. Subject FA TRR in the HCP-TR also tends to be high, but the values vary more across bundles with a range of 0.57 ±0.24 to 0.85 ±0.12 and a median across bundles of 0.73. We can see that subject TRR is lower than profile TRR (Figure 2). This trend is consistent for MD (Fig. S5) as well as for another dataset (Fig. 3**C**).

### Test-retest reliability of tractometry in different implementations, datasets, and tractography methods

We compared TRR across datasets and implementations. In both datasets, we found high TRR in the results of tractography and bundle recognition: wDSC was larger than 0.7 for all but one bundle (Fig. 3**A**): the delineation of the anterior forceps (FA bundle) seems relatively unreliable using pyAFQ in the UW-PREK dataset (using the FA scalar, pyAFQ subject TRR is only 0.37 ±0.28 compared to mAFQ’s 0.84 ±0.10). We found overall high profile TRR that did not always translate to high subject TRR (Fig. **3B-G**). For example, for FA in UW-PREK, median profile TRRs are 0.75 for pyAFQ and 0.77 for mAFQ while median subject TRRs are 0.70 for pyAFQ and 0.75 for mAFQ. Note that profile and subject TRR have different denominators (for example, subjects that have similar mean profiles to each other would have low subject TRR, even if the profiles are reliable, because it is harder to distinguish between subjects in this case). mAFQ is one of the most popular software pipelines currently available for tractometry analysis, so it provides an important point for comparison. In comparing different software implementations, we found that mAFQ has higher subject TRR relative to pyAFQ in the UW-PREK dataset, when TRR is relatively low for pyAFQ (see the FA bundle, CST L, and ATR L in Fig. 3**C**). On the other hand, in the HCP-TR dataset pyAFQ we used the RTP pipeline (42,43), which is an extension of mAFQ, and found that pyAFQ tends to have slightly higher profile TRR than RTP for MD, but slightly lower profile TRR for FA (Fig. 3**D**). The pyAFQ and RTP subject TRR are highly comparable (Fig. 3**E**). In FA, the median pyAFQ subject TRR for FA is 0.76 while the median RTP subject TRR is 0.74. Comparing different ODF models in pyAFQ, we found that the DKI and CSD ODF models have highly similar TRR, both at the level of wDSC (Fig. 3**A**), as well as at the level of profile and subject TRR (Fig. **3F-G**).

### Robustness: comparison between distinct tractography models and bundles recognition algorithms

To assess the robustness of tractometry results to different models and algorithms, we used the same measures that were used to calculate TRR.

#### Tractometry results can be robust to differences in ODF models used in tractography

We compared two algorithms: tractography using DKI- and CSD-derived ODFs. The weighted Dice similarity coefficient (wDSC) for this comparison can be rather high in some cases (e.g., the uncinate and corticospinal tracts, Figure 4A), but produce results that appear very different for some bundles, such as the arcuate and superior longitudinal fasciculi (ARC and SLF) (see also Figure 4D). Despite these discrepancies, profile and subject robustness are high for most bundles (median FA of 0.77 and 0.75, respectively) (Figure **4B,C**). In contrast to the results found in TRR, MD subject robustness is consistently higher than FA subject robustness. The two bundles with the most marked differences between the two ODF models are the SLF and ARC (Figure 4**D**). These bundles have low wDSC and profile robustness, yet their subject robustness remains remarkably high (In FA, 0.75 ±0.17 for ARC R and 0.88 ±0.09 for SLF R) (Figure 4**C**). These differences are partially explained due to the fact that there are systematic biases in the sampling of white matter by bundles generated with these two ODF models, as demonstrated by the non-zero adjusted contrast index profile (ACIP) between the two models (Figure 4E).

**Fig. 4.**
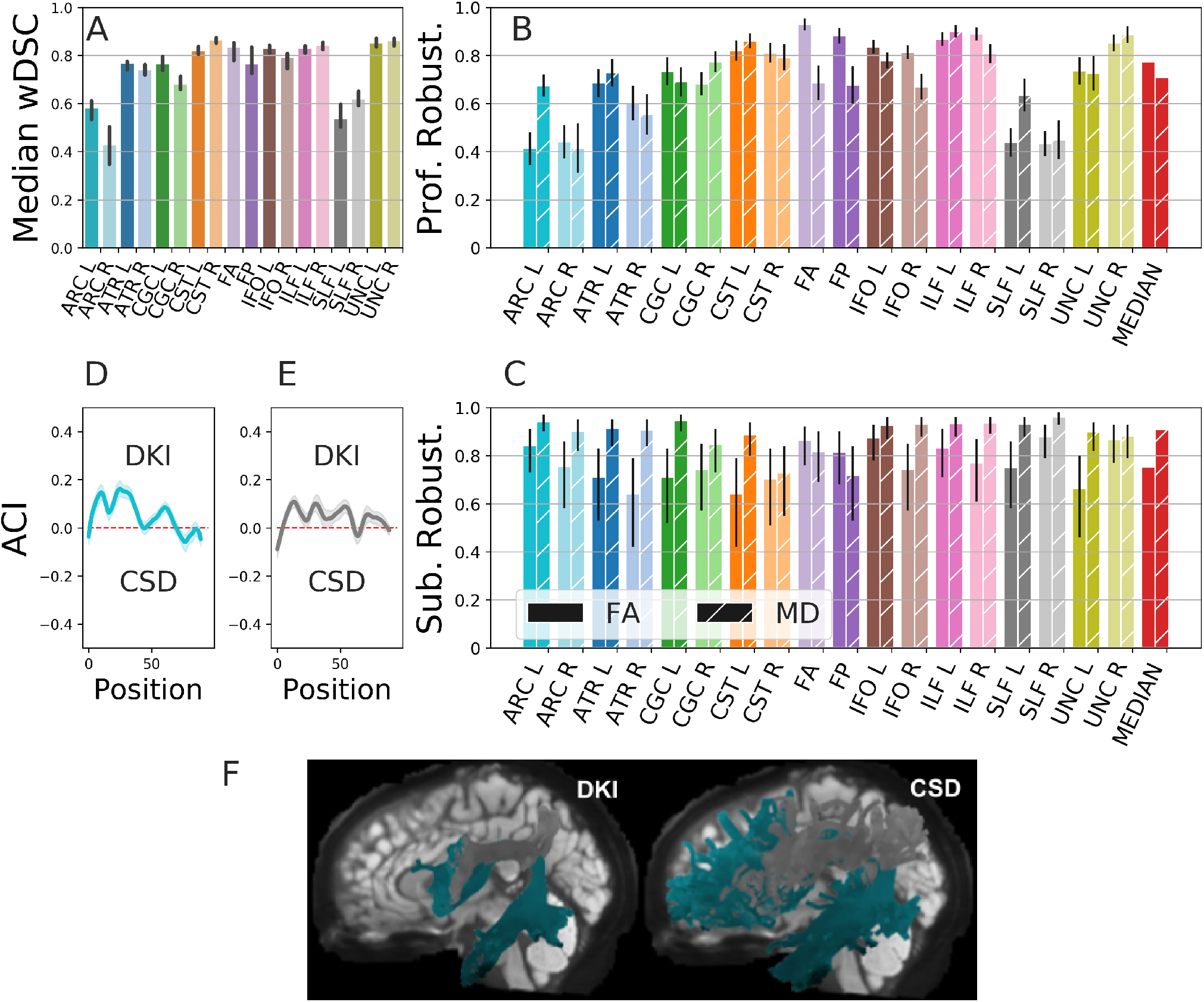
ODF model robustness. We compared DKI- and CSD-derived tractography. Colors encode bundle information as in Figures 1 and 2. Textured hatching encodes FA/MD information. **A** wDSC robustness. **B** Profile robustness. **C** Subject robustness. Error bars represent 95%confidence interval. **D**, **E** Adjusted contrast index profile (ACIP) between ARC L and SLF L tract profiles of each algorithm. Positive ACI indicates DKI found a higher value of FA than CSD at that node. The 95%confidence interval on the mean is shaded. **F** Tractography and bundle recognition results for ARC L and SLF L respectively for one example subject.

#### Most white matter bundles are highly robust across bundle recognition methods

We compared bundle recognition with the same tractography results using two different approaches: the default waypoint ROI approach (9), and an alternative approach (RecoBundles) that uses atlas templates in the space of the streamlines (44). Between these algorithms, wDSC is around or above 0.6 for all but one bundle, ILF R (Figure 5). There is an asymmetry in the ILF atlas bundle(7), which results in discrepancies between ILF R recognized with way-point ROIs and with RecoBundles. Despite this bundle, we find high robustness overall. For MD, the first quartile subject robustness is 0.82 (Figure 5C, D).

**Fig. 5.**
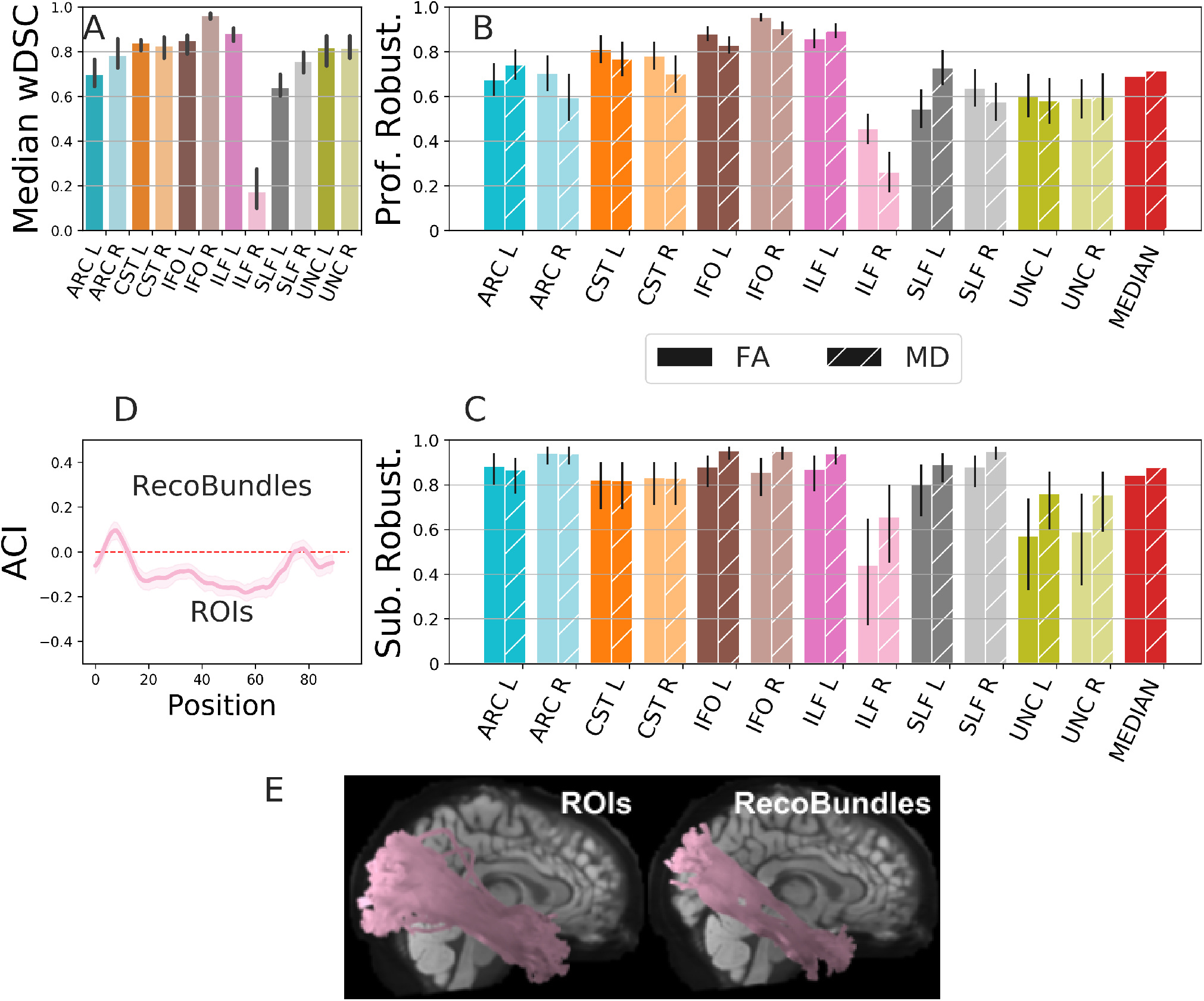
Recognition algorithm robustness. **A** wDSC. **B** Profile robustness. **C** Subject robustness. Error bars show the 95% confidence interval. **D** The ILF R FA ACIP, where positive ACI indicates RecoBundles found a higher value of FA than the waypoint ROIs approach at that node. **E** shows the ILF R found by each algorithm for an example subject.

#### Tractometry results are robust to differences in software implementation

Overall, we found that robustness of tractometry across these different software implementations is high in most white matter bundles. In the mAFQ/pyAFQ comparison, most bundles have a wDSC around or above 0.8, except the two callosal bundles (FA bundle and FP), which have a much lower overlap (Fig. 6A). Consistent with this pattern, profile and subject robustness is also overall rather high (Fig. 6**B**, C). The median values across bundles are 0.71 and 0.77 for FA profile and subject robustness, respectively.

**Fig. 6.**
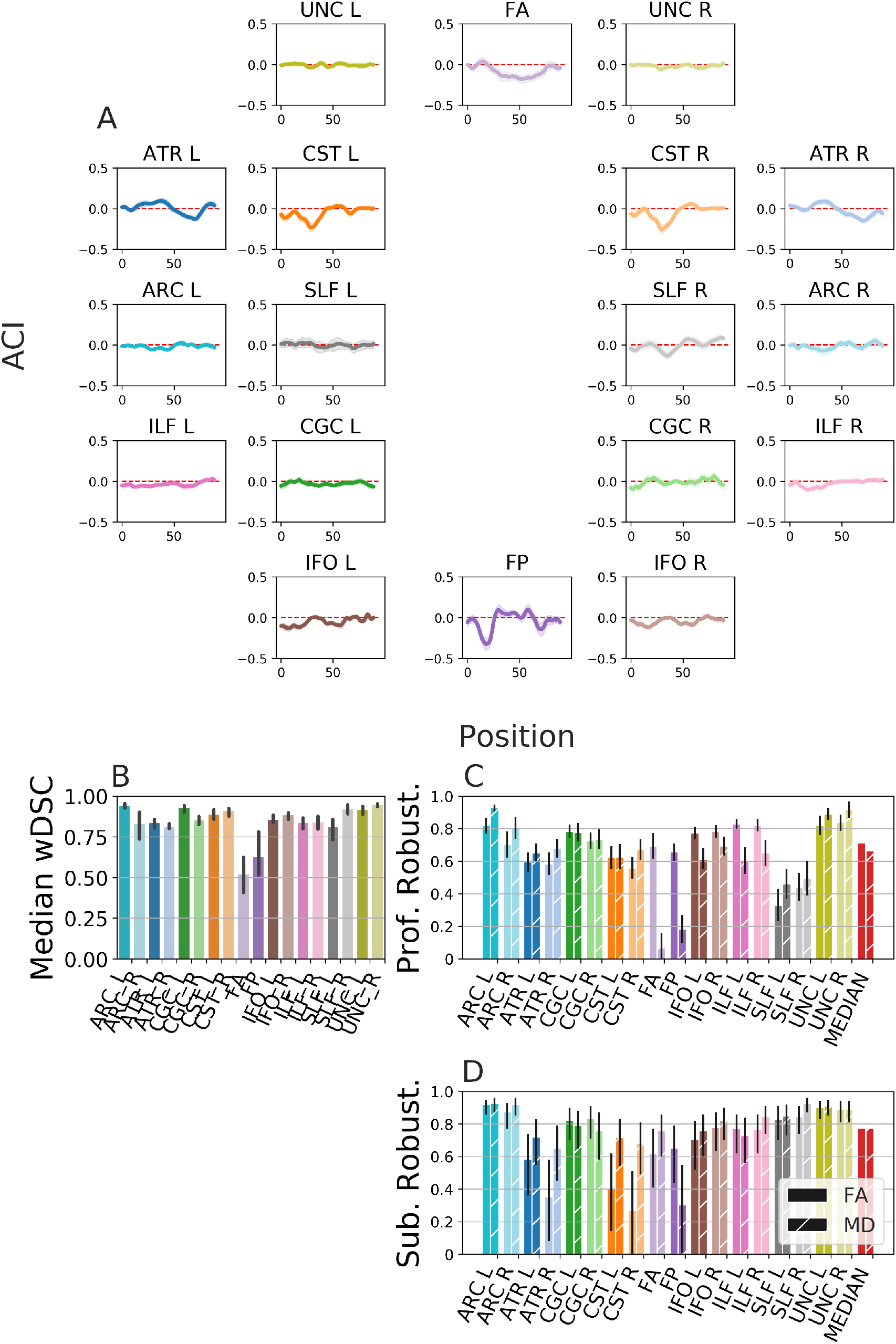
Robustness between pyAFQ and mAFQ on UW-PREK session # 1 data. **A** ACIP between the FA tract profiles from UW-PREK using pyAFQ and mAFQ. Positive ACI indicates pyAFQ found a higher value than mAFQ at that node. The 95% confidence interval on the mean is shaded. Robustness in wDSC (**B**) bundle profiles (**C**) and across subjects (**D**). Error bars show the 95% confidence interval.

For some bundles, like the right and left uncinate, there is large agreement between pyAFQ and mAFQ (for subject FA: UNC L *ρ* = 0.90 ±0.07, UNC R *ρ* = 0.89 ±0.08). However, the callosal bundles have particularly low mean diffusivity (MD) profile robustness (Fig. 6B) (0.07 ±0.09 for FP, 0.18 ±0.09 for FA).

The robustness of tractometry to the differences between the pyAFQ and mAFQ implementation depends on the bundle, scalar, and reliability metric. In addition, for many bundles, the ACIP between mAFQ and pyAFQ results is very close to 0, indicating no systematic differences (Fig. 6D). In some bundles – the corticospinal tract (CST) and the anterior thalamic radiations (ATR) – there are small systematic differences between mAFQ and pyAFQ. In the Forceps Posterior (FP), pyAFQ consistently finds smaller FA values than mAFQ in a section on the left side. Notice that the forceps anterior has an ACIP that deviates only slightly from 0, even though the forceps recognitions did not have as much overlap as other bundle recognitions (see Fig. 6A).

## Discussion

Previous work has called into question the the reliability of neuroimaging analysis (e.g., (25,45,46)). We assessed the reliability of a specific approach, tractometry, which is grounded in decades of anatomical knowledge, and we demonstrate that this approach is reproducible, reliable and robust. A tractometry analysis typically combines the outputs of tractography with diffusion reconstruction at the level of the individual voxels within each bundle. One of the major challenges facing researchers who use tractometry is that there are many ways to analyze diffusion data, including different models of diffusion at the level of individual voxels; techniques to connect voxels through tractography; and approaches to classify tractography results into major white matter bundles. Here, we analyzed the reliability of tractometry analysis at several different levels. We analyzed both test-retest reliability of tractometry results and their robustness to changes in analytic details, such as choice of tractography method, bundle recognition algorithm, and software implementation (Fig 6).

### Test-retest reliability of tractometry

Test-retest reliability (TRR) of tractometry is usually rather high, comparable in some tracts and measurements to the TRR of the measurement. In comparing the HCP-TR analysis and UW-PREK analysis, we note that higher measurement reliability goes hand in hand with tractometry reliability.

In terms of the anatomical definitions of the bundles, quantified as the TRR wDSC, we find reliable results in both datasets and with both software implementations and both tractography methods that we tested. With pyAFQ we found a relatively low TRR in the frontal callosal bundle (FA bundle) in the UW-PREK dataset. This could be due to the sensitivity of the definition of this bundle to susceptibility distortion artifacts in the frontal poles of the two hemispheres. This low TRR was not found with mAFQ, suggesting that this low TRR is not a necessary feature of the analysis, and is a potential avenue for improvement to pyAFQ. While the two implementations were created by teams with partial overlap and despite the fact that pyAFQ implementation drew both inspiration as well as specific implementation details from mAFQ, many details of implementation still differ substantially. For example, the implementations of tractography algorithms are quite different – pyAFQ relies on DIPY (28) for its tractography, while mAFQ uses implementations provided in Vistasoft (47). The two pipelines also use different registration algorithms, with pyAFQ relying on the SyN algorithm (33), while mAFQ relies on registration methods implemented as part of the Statistical Parametric Mapping (SPM) software (48). These differences may explain the discrepancies observed.

We also find that TRR is high at the level of profiles within subjects and mean tract profiles across subjects. This is generally observed in both datasets that we examined, and using different analysis methods and software implementations. For the UW-PREK dataset, subject TRR tends to be higher in mAFQ than in pyAFQ. On the other hand, for the HCP-TR dataset, pyAFQ subject TRR tends to be higher than that obtained with RTP, which is a fork and extension of mAFQ (42,43). Generally, TRR of FA profiles and also TRR of mean FA across subjects tend to be higher than those of MD. This could be because the assessment of MD is more sensitive to partial volume effects. In contrast to FA, MD is also not bounded, which means that extreme values at the boundaries of tissue types can have a substantial effect on TRR.

### Robustness of tractometry

As highlighted in the recent work by Botvinik-Nezer *et al* (25) and in parallel by Schilling *et al* (45), inferences from even a single dataset can vary significantly, depending on the decisions and analysis pipelines that are used. The analysis approaches used in tractometry embody many assumptions made at the different stages of analysis: the model of the signal in each individual voxel, the manner in which streamlines are generated in tractography, the definition of bundles, and the extraction of tract profiles. While TRR is important, it does not guard against systematic errors in the analysis approach. One way to test model assumptions and software failures is to create ground truth data against which different methods and implementations can be tested (13,49,50). However, this approach also relies on certain assumptions about the mechanisms that generate the data that is considered ground truth, making this approach more straightforward for some methods than others. Here, we instead assessed the robustness of tractometry results to perturbations of analytic components, focusing on the modelling of ODFs in individual voxels and the approach taken to bundle recognition.

#### Subject robustness remains high despite differences in the spatial extent of bundles

We replicated previous findings that the definition of major bundles can vary in terms of their spatial extent (quantified via wDSC) (13,37,40,45), depending on the software implementation or the ODF model used. As we show, low wDSC robustness often corresponds to low profile robustness, and vice versa (Fig **6B,C**, Fig 4A,B, and Fig 5A,B). That is, when two algorithms detect bundles with small spatial overlap, the shape of the resulting tract profiles are also different from each other. However, low wDSC and profile robustness does not always translate to low subject robustness. Algorithms can detect bundles with low spatial overlap and of different shapes yet still agree on the ordering of the mean of the profiles, i.e., which subjects have high or low FA in a given bundle. A clear example of this is the SLF and ARC in Fig 4 (wDSC and profile robustness are low, yet subject robustness is very high). This suggests that tractometry can overcome failures in precise delineation of the major bundles by averaging tissue properties within the core of the white matter. Conversely, important details that are sensitive to these choices may be missed when averaging along the length of the tracts. Moreover, this may also reflect biases in the measurement that cannot be overcome at either stage of the analysis: tractography or bundle recognition.

Our high subject-level robustness results (Fig 6**C**, Fig 4C, and Fig 5C) dovetail with the results of a recently-published study that used tractometry in a sample of 45 participants (51), and found high subject-level correlations between the mean tract values of FA and MD for two different pipelines: deterministic tractography using the diffusion tensor model (DTI) as the ODF model (essentially identical to a pipeline used in our supplementary analysis, described in “DTI Configuration”), and probabilistic tractography using CSD as the ODF model. Consistent with our results on the HCP-TR dataset, slightly higher subject robustness was found for MD than for FA.

#### Exceptions & Limitations

High profile robustness did not always imply high subject robustness (e.g., the FP in Fig 4 has high profile robustness, but low subject robustness). This suggests that there are other sources of between-subject variance that do not correspond directly to profile robustness within an individual.

There are still significant challenges to robustness that arise from the way in which the major bundles are defined. This problem was highlighted in recent work that demonstrated that different researchers use different criteria to define bundles of streamlines that represent the same tract (45). In our case, this challenge is represented by the relatively low robustness between the waypoint ROI algorithm for bundle definition and the RecoBundles algorithm. In this comparison, the wDSC exceeds 0.8 in only one bundle and is below 0.4 in two cases. While both algorithms identify a bundle of streamlines that represents the right ILF, this bundle differs substantially between the two algorithms. Even so, profile and subject robustness can still be rather high, even in some cases in which rather middling overlap is found between the anatomical extent of the bundles. This challenge highlights the need for more precise definitions of the models of brain tracts that are derived from dMRI, but also highlights the need for clear, automated and reproducible software to perform bundle recognition.

In addition to decisions about analysis approach, which may be theoretically motivated, software implementations may contain systematic errors in executing the different steps and different software may be prone to different kinds of failure modes. Since other software implementations (9,42) of the AFQ approach have been in widespread use in multiple different datasets and research settings, we also compared the results across different software implementations (Fig. 6). While there are some systematic differences between implementations, tractometry is overall quite robust to differences between software implementations.

Another important limitation of this work is that we have only analyzed samples of healthy individuals. Where brains are severely deformed (e.g., in TBI, brain tumors and so forth), particular care would be needed to check the results of bundle recognition, and separate considerations would be needed in order to reach conclusions about the reliability of the inferences made.

### Computational reproducibility via open-source software

Reproducibility is a bedrock of science, but achieving full computational reproducibility is a high bar that requires access to the software, data and computational environment that a researcher uses (22). One of the goals of pyAFQ is to provide a platform for reproducible tractometry. It is embedded in an ecosystem of tools for reproducible neuroimaging and is extensible. This is shown in Fig. S6 and Fig S2 and is further discussed in “Supplementary Discussion of pyAFQ”. Results from the present article and supplements can be reproduced using a set of Jupyter notebooks provided here: https://github.com/36000/Tractometry_TRR_and_robustness. After installing the version of pyAFQ that we used (0.6), reproduction should be straight-forward on standard operating systems and architectures, or in cloud computing systems (see code and Supplementary Methods). In the UW-PREK dataset, we shared the tract profiles and we provide web-based visualizations using a tool that previously developed for transparent data sharing of tractometry data (52): https://yeatmanlab.github.io/UW_PREK_pyAFQ_pre_browser and https://yeatmanlab.github.io/UW_PREK_pyAFQ_post_browser.

The HCP-TR dataset is relatively straightforward for others to access in its preprocessed form through the HCP, and because the study IDs can be openly shared in our code, anyone with such access should be able to reproduce the figures in full. Using these resources, it should be possible to re-execute our workflows and replicate most of our results (53). For example, if other researchers would be interested in comparing our TRR results to another tractometry pipeline (e.g., TRACULA (11), another popular tractometry pipeline) or another bundle recognition algorithm (e.g., TractSeg (54), which uses a neural network to recognize bundles, or Classifyber (55) which uses a linear classifier), they could do so with the HCP-TR dataset, inspired by our scripts, and the visualization tools in the pyAFQ software.

### Future Work

There are many aspects of reliability that could be further explored. We explored robustness with respect to ODF models and bundle recognition algorithms; robustness could also be explored with respect to: data acquisition parameters within the same subject; preprocessing methods; profile extraction method (for example, comparing our current approach with the BUndle ANalytics (BUAN) (56)); and the effects of profile realignment on tract profile reliability (57). Another possibility for teasing apart measurement and tractography effects would be to test profile TRR using the streamline of one scan on the results of the second scan (by registering the streamline themselves, to avoid data interpolation in volume registration). This could tease apart the effects of tractography from the voxel-level models of tissue properties, because it is not necessary that these would be sensitive to the same constraints (e.g., different sensitivity to noise). The methods we demonstrate and resources we provide in this paper should be useful for anyone wishing to further explore reliability in tractometry.

## ACKNOWLEDGEMENTS

This work was supported through grant 1RF1MH121868-01 from the National Institute of Mental Health/The BRAIN Initiative, through grant 5R01EB027585-02 to Eleftherios Garyfallidis (Indiana University) from the National Institute of Biomedical Imaging and Bioengineering and through Azure Cloud Computing Credits for Research & Teaching provided through University of Washington Research Computing and the University of Washington eScience Institute. We are also grateful for support from the Gordon & Betty Moore Foundation and the Alfred P. Sloan Foundation to the University of Washington eScience Institute Data Science Environment, as well as support from the Washington Research Foundation to the eScience Institute and to the University of Washington Institute for Neuroengineering. Thanks to Andreas Neef for feedback on the pyAFQ software. Data were provided in part by the Human Connectome Project, WU-Minn Consortium (Principal Investigators: David Van Essen and Kamil Ugurbil; 1U54MH091657) funded by the 16 NIH Institutes and Centers that support the NIH Blueprint for Neuroscience Research; and by the McDonnell Center for Systems Neuroscience at Washington University.

## Supplementary Methods

### Automated Fiber Quantification in Python (pyAFQ)

Inspired by a previous MATLAB implementation (9), We developed a software library that automates dMRI-based tractometry analysis. The library is called pyAFQ (Python Automated Fiber Quantification), and it is implemented as open-source software here: https://github.com/yeatmanlab/pyAFQ. The software is developed under the permissive OSI-approved BSD license. It allows users to specify the methods and parameters they want to use for tractometry. pyAFQ uses many components of the scientific Python ecosystem (58). In particular, it relies heavily on implementations of algorithms for diffusion reconstruction, orientation determination, tractography and image registration implemented in Diffusion Imaging in Python (DIPY), an open-source, Python library for computational neuroanatomy (28). The pyAFQ software implements extensive documentation with Sphinx (59), including a gallery of executable examples, implemented using Sphinx Gallery (60). Unit testing is implemented using pytest, with continuous integration implemented to test proposed changes to the library, as well as longer nightly tests that check that pipelines of operations are not adversely affected by changes that are introduced in developing the software. pyAFQ’s test suite uses the HARDI data collected for (16), CFIN (61), and data from the Human Connectome Project. pyAFQ can be parallelized across subjects and sessions using dask (62). The analysis performed in this paper primarily used pyAFQ run using Cloudknot (63) on Amazon Web Services (AWS).

There are many ways to analyze dMRI data and to estimate tractomery-based tract-profiles. For example, many different models are used to determine the directions of tracking within each voxel and to connect different voxels with a variety of tractography algorithms. Similarly, different models can be used to determine the tissue properties within a voxel. However, it is hard to determine which methods to use, because different methods may be appropriate for different datasets, depending on their characteristics: the measurements conducted, the signal to noise ratio (SNR) of the data and so forth. Software to support analysis of a variety of datasets should make it easy to use many different methods and to compare results between methods. All of the choices the user can make in each of the steps of pyAFQ are delineated below and summarized in Fig. S2. The software implements a library with an object-oriented application programming interface (API), as well as a command-line interface (CLI). Using pyAFQ’s API, pyAFQ can be run with only a few lines of code. The API is also flexible, giving the user the ability to choose which algorithms and parameters to use. For users unfamiliar with python, pyAFQ has a command line interface (CLI) which uses a configuration file written in TOML (64). pyAFQ also has a Boutiques configuration file and can be executed using Boutiques (65).

#### Locating and mapping data (BIDS)

The first step in analysis is to find the files that the software will use. pyAFQ relies on pyBIDS (66,67) to query data that is provided in the BIDS format (68). It looks for dMRI, b-value, and b-vector files stored in standard formats (see https://yeatmanlab.github.io/pyAFQ/usage/data.html for details). Additionally, the user can provide files from other processing pipelines to be used as a brain mask during registration or as start or stop masks during tractography, as well as completed tractography results. We typically use the Nibabel software library to interact with neuroimaging files (69). Following the BIDS standard, the outputs of pyAFQ are put in the BIDS derivatives folder, in a pipeline directory labelled as “afq”. The derivative BIDS format follows as much as possible the draft implementation of the BIDS derivatives for dMRI data.

#### Tractography

There are several methods for computational tractography. The pyAFQ software exposes many of these as options. It allows users to choose from multiple fiber orientation distribution functions (70) that determine the direction of tracking in each step of the process: based on Diffusion Tensor Imaging (DTI) (71,72), Diffusion Kurtosis Imaging (DKI) (73), Constrained Spherical Deconvolution (CSD) (74,75), and Multi-Shell Multi-Tissue Constrained Spherical Deconvolution (MSMT-CSD) (76). Deterministic and probabilistic tractography algorithms can be used and stopping criteria can be implemented for particle filtering tractography, using the continuous map criterion (77) or anatomically-constrained tractography (78). The default tractography setting uses DTI, deterministic direction finding, a max turning angle per step of 30°, one seed per voxel, and retains only streamlines between 10 and 1000mm long. Many of our tractography defaults are inspired by the results of (79) and (9). The default seed and stop masks are created by thresholding FA at 0.2. All of these parameters can be customized using pyAFQ’s API or CLI.

#### Template registration

The user can specify their own template and subject image to register, however pyAFQ also provides four builtin options: register subject non-diffusion weighted image (also known as b0) to the Montreal Neurological Institute (MNI) T2 template (29,30); register subject FA to a group mean fractional anisotropy (FA) template from the UK Biobank (80,81); register a subject’s anisotropic power map (APM) (31,32) to the MNI T1 template; and register subject streamlines to the 16 bundles human connectome project (HCP) atlas (7) using streamline registration (SLR) (82). The first three of these builtin techniques use the nonlinear Symmetric Diffeomorphic Registration (SyN) (33) after an optional linear preregistration, both implemented in DIPY. pyAFQ uses Templateflow (83) to get MNI T1/T2 templates for registration. The default registration behavior is to consider all b-values under 50 to be b0, mask the subject’s APM using DIPY’s median_otsu image recognition algorithm (84) on the subject b0, and register the masked power map to the masked MNI T1 template. Per default, we chose to use the APM for registration based on previous findings that show this is a good choice (85) and based on our own experience. All of these parameters can be customized using pyAFQ’s API and CLI.

#### Bundle recognition and cleaning

To identify the streamlines that best represent a particular anatomical pathway, we perform bundle recognition. The default behavior is to perform the initial classification using probability maps, and then segment with waypoint ROIs defined in (86), then filter the classified streamlines by their termination locations, using the AAL atlas (87), where streamlines must be within 4mm of the expected endpoint region. Waypoint ROIs are moved into the subject space and then patched up using the Quickhull Algorithm (88). There is also an option, turned off by default, to clip streamline edges at the ROIs (86).

In addition to the waypoint-based recognition described above, pyAFQ also allows the user to choose to use a streamline atlas based bundle recognition method, called RecoBundles (44). Parameters for either algorithm can be customized using pyAFQ’s API and CLI.

After recognition, cleaning is performed based on the Mahalanobis distance of each streamline from the mean in each node. This process was originally described in (9). By default, pyAFQ resamples streamlines to 100 points (nodes) and performs 5 rounds of cleaning with a distance threshold of 5 standard deviations from the mean of the node coordinates at each point, and a length threshold of 4 standard deviations from the mean length. Cleaning is also stopped if a bundle has less than 20 streamlines. All of these parameters can be customized using pyAFQ’s API and CLI.

#### Tract Profile Extraction

After cleaning, pyAFQ computes and visualizes tract profiles. The mean profile (called a “tract profile”) is calculated using the same Mahalanobis distance-based weighting strategy as in Yeatman et al. (9), implemented in DIPY. Visualization can be performed using one of two backends: fury (89) or plotly (90), which create either animated gifs or interactive html files respectively. Visualizations are created for the whole brain tractometry and for each individual bundle.

### Data

We measured the reliability of tractometry using two datasets with contrasting characteristics.

#### Human Connectome Project (HCP-TR)

The WU-Minn Human Connectome Project (HCP) (91) includes measurements of diffusion MRI data from almost all of the 1,200 participants. Here, we focus our analysis on a subset of these subjects for which test-retest data are available. We refer to this data as HCP-TR. This dataset contains dMRI data from 44 individuals. This represents a relatively high-quality, high-resolution dataset, with multiple diffusion directions and multiple b-values. The acquisition parameters of HCP-TR are described in detail elsewhere (36). We used data that had been preprocessed through the HCP pipelines, as provided through the AWS Open Data program (https://registry.opendata.aws/hcp-openaccess/).

#### University of Washington Pre-K (UW-PREK)

Two measurements were conducted in each participant 1 day apart. These were acquired with 32 directions, b=1,500 s/mm^2^, 2 mm^3^ isotropic resolution, TR/TE=7200/83 msec. Data were preprocessed using FSL for eddy current, motion correction, and susceptibility distortion correction. Analysis using the mAFQ was conducted as previously described (9). We converted UW-PREK to BIDS format (68) for input into pyAFQ’s API.

We attempted to configure pyAFQ to most closely match the mAFQ configuration. We used robust estimation of tensors by outlier rejection (RESTORE) (92) to fit the DTI model. In tractography, we used 160,000 seeds randomly distributed wherever DTI FA is higher than 0.3. We used only 1 round of cleaning. We ran this on both the UW-PREK pre and post sessions, and compared its reproducibility to the results on the same datasets with mAFQ. We also compared the robustness of the results between the pyAFQ and mAFQ algorithms on the pre-session data only.

### Configurations

For all configurations, we used the Freesurfer brain segmentation provided by HCP to calculate a permissive brain mask, with all portions of the image not labelled as 0, considered part of the brain. The brain mask is used when fitting the ODF models. We compared the TRR of each configuration, as well as the robustness of the results across configurations. We also compared the TRR of these configurations to the TRR of results published by Lerma-Usabiaga and colleagues (43), denoted RTP.

#### DTI Configuration

In addition to the three configurations enumerated in the present paper, we processed HCP-TR with a fourth configuration. We used only measurements with b-values between 990 and 1010 s/mm^2^. We used DTI as the ODF model for tractography and profile extraction. We compared this configuration to RTP in **3D,E**. We also analysed DTI for robustness and found its results to be nearly identical to DKI.

#### RecoBundles Configuration

One of the configurations we ran on the HCP-TR data used RecoBundles (8). pyAFQ provides programmatic access to two atlases, one being the full 80 bundles human connectome project (HCP) atlas (7), and other being a 16 bundle subset of that atlas. We ran RecoBundles on HCP-TR using the full 80 bundles atlas. We use the following RecoBundles parameter configuration: a model cluster threshold of 1.25, a reduction threshold of 25, no refinement, a pruning threshold of 12, local streamline-based linear registration on with an asymmetric metric. We used this configuration for all 80 bundles. Multi-shell data and the DKI ODF model were used. We used nonlinear symmetric diffeomorphic registration and a brain mask based on the HCP-provided segmentation.

#### RTP

As a point of comparison, we used an open dataset of HCP-TR derivatives that was published by Lerma-Usabiaga and colleagues (43). They processed HCP-TR using the Reproducible Tract Profiles (RTP) pipeline (42). This pipeline is a full end-to-end pipeline and system for deployment of analysis that receives as input raw MRI data as acquired on the scanner. While it applies different preprocessing steps and uses different tractography algorithms than mAFQ, relying on MRTRIX for many of these steps (93), the bundle recognition steps closely resemble the ones used in mAFQ, relying on functions that stem from the same MATLAB codebase as mAFQ. The end result of RTP are tract profiles in an easy-to-use and data-science ready JSON format. We denote their results as RTP and compare them to the HCP-TR results computed with pyAFQ.

### Measures of reliability

pyAFQ gives the user the choice of which underlying algorithms to use when performing tractometry, as shown in Fig. S2. We use this feature of pyAFQ to run multiple analyses on HCP-TR and UW-PREK, which both have test-retest data. The analyses we selected represent only a small subset of the possible configurations of pyAFQ. However, because the software is freely available and easily configurable with the API or CLI, it would be straightforward to test other analyses. To compare the results on test-retest data (TRR) and compare results across analyses (robustness), we use four different measures of reliability. Each one of these measures emphasizes different aspects of reliability.

#### Weighted Dice similarity coefficient (wDSC)

The anatomical reliability of bundle recognition solutions is assessed by comparing their spatial overlap in the white matter volume. First, for every voxel in the white matter, we count the number of streamlines that pass through that voxel for a given bundle, then divide by the total number of streamlines in that bundle. This creates what we call a streamline density map (28). We could compare streamline density maps using a Dice similarity coefficient (94), but that would require applying a threshold to the density maps, and could give a few streamlines a large influence on the calculation. Instead, we use the weighted Dice similarity coefficient (wDSC) (37):

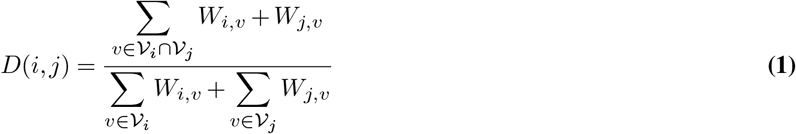

where *ν* is a voxel index, *W_i,ν_* is the streamline density for a bundle *i* in voxel *ν* and *ν*’ are voxels where the two bundles *i* and *j* intersect. wDSC provides a measure of the reliability in the spatial extent of bundles, in a manner that is independent from the assessment of tract profiles.

#### Adjusted contrast index profile (ACIP)

We use an adjusted contrast index to directly compare the values of individual nodes in the tract profiles in different measurements. For two values (*V*_1_, *V*_2_) in different profiles, the adjusted contrast index (ACI) is calculated using Eq (2).

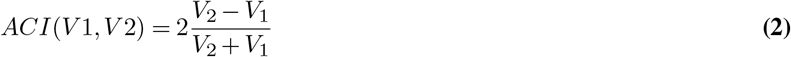

We multiply by 2 to make the contrast index have comparable values to fractional difference. In contrast to fractional difference, however, the ACI does not require one of the variables to be a reference, and *ACI*(*V*1,*V*2) =−*ACI*(*V*2,*V*1). Calculating and then plotting the ACI for each point between two profiles highlights the differences between profiles, producing the adjusted contrast index profile (ACIP). ACIP emphasizes discrepancies in estimates along the length of the tract in a manner that does not depend on the scale of the measurement (e.g., the different scales of FA and MD).

## Supplementary Discussion of pyAFQ

### pyAFQ is embedded in an ecosystem of tools for reproducible neuroimaging

The wider ecosystem of tools and standards surrounding pyAFQ is shown in Fig. S6. Each tool has its own place in the ecosystem. We rely heavily on implementations of dMRI analysis algorithms implemented in DIPY (28). Reproducibility and interoperability are also facilitated by relying on the BIDS format (68) and the pyBIDS software (66,67). Requiring a BIDS-like input makes integration with other software in the ecosystem easier. For example, it is fairly straightforward to use the outputs of BIDS-compatible preprocessing pipelines, such as qsiprep (95), as inputs to pyAFQ. Furthermore, the modularity of the pyAFQ pipeline means that outputs of other tractography software (e.g., MRTRIX (96)) can be used as inputs to bundle recognition, with BIDS filters as the metadata that allows finding and incorporating through the right data.

Cloud-based processing is going to be more important as large datasets are processed. pyAFQ does not depend on proprietary software and can be scaled to large datasets using cloud computing platforms. In this paper, we used Cloudknot (63) to scale pyAFQ across subjects and methods on AWS. However, because pyAFQ is a Python package, it can easily be run on any cloud computing platform. Computing in the public cloud also supports reproducible research, as computations conducted on the public cloud are perfectly portable to other users of the software. Our software is written with that in mind, including functions that know how to easily access datasets that are already stored in the cloud (e.g., HCP and Healthy Brain Network (97) datasets). We know that one of the most important ways in which users can diagnose whether processing worked as expected is by visually inspecting the results. Thus, we provide several different visualization methods, relying on the VTK-derived FURY library, or on browser-friendly visualizations with Plotly. pyAFQ outputs are also fully compatible with AFQ-Browser, a browser-based tool for interactive visualization and exploration of tractometry results (52).

Finally, beyond visualization and summary of the results, and tools for analysis of reliability presented in this work, pyAFQ does not provide a substantial set of tools for statistical analysis of tractometry results. Instead, the outputs of pyAFQ are provided as “tidy” CSV tables (27). This means that it is compatible as inputs to the AFQ Insight tool for statistical analysis (20), but also amenable to many other statistical analysis approaches. This output should facilitate interdisciplinary use of dMRI data, as it is provided in a format that is widely used in statistics and machine learning.

### pyAFQ is extensible

In general, variability in results would be reduced with a standard pipeline that could be used across all studies and datasets. However, as noted by Lindquist, “studies tend to be too varied for one pipeline to always be appropriate” (98). This is particularly true as new measurement techniques, new processing methods and new analysis approaches for dMRI are evolving. Therefore, the pyAFQ pipeline was designed to be flexible, making it easier to reproduce results, while providing researchers with many choices for the appropriate analysis, depending on their data and questions. pyAFQ allows the user to make many decisions (Fig S2), and all of those decisions can be encoded in a configuration file. That configuration file can be used to reproduce the same analysis pipeline given the same version of pyAFQ is used. By providing the configuration file or the arguments passed to the main API, one can clearly satisfy the requirement for a re-executable workflow outlined in (53).

To extend to new bundles, pyAFQ allows users to define new queries that recognize bundles that are not part of the set of 18 detected by the original mAFQ software. For a simple example, we use a set of alternative waypoint ROIs to detect different portions of the corpus callosum (99) (Fig S7**A**). These alternative ROIs are included in pyAFQ but not used by default. In more complicated example, another set of ROIs is used to recognize the location of the optic radiations (OR; Fig S7). Because these are relatively small and winding, their delineation requires additional components: it requires several waypoint ROIs used not only as inclusion criteria, but also as exclusion criteria, and it requires delineation of endpoints in the cortex that are not part of the AAL atlas, which is used in the standard set of bundles. It also requires oversampling of streamlines, so in order to obtain a proper definition of the OR, tractography is configured to use 125 seeds per voxel (instead of the default 8). All of these components can be integrated into calls to the software API, without needing to change any of its internals. This includes any custom waypoint ROIs, inclusive or exclusive, as well as probability maps, endpoint locations, and whether the bundle crosses the midline.

## Supplementary Figures and Tables

**Fig. S1.**
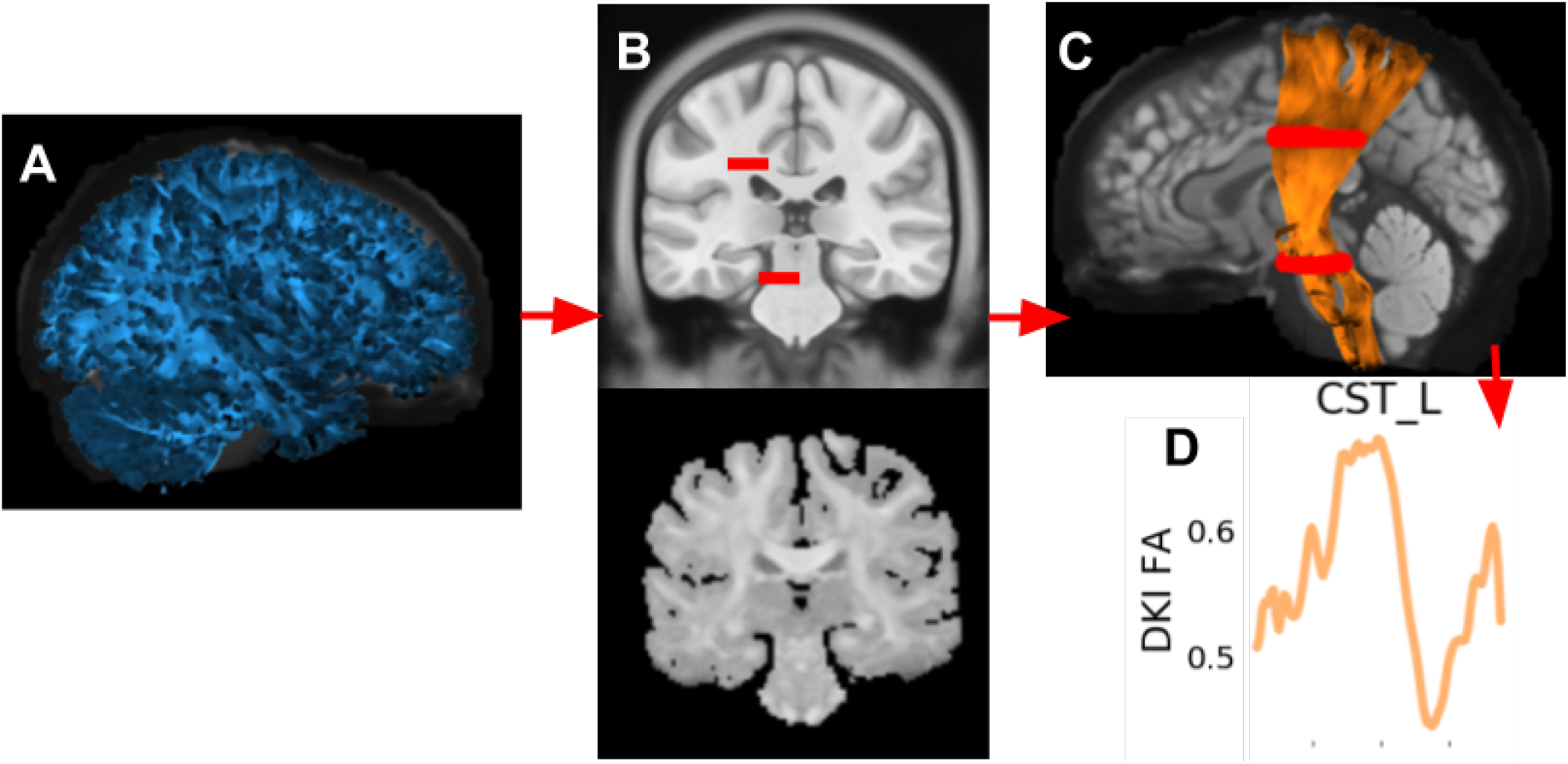
The stages of tractometry. **A** Computational tractography generates streamlines estimating the trajectories of white matter connections. **B** An anatomical template is registered to each subjects individual brain. Here, in a mid-coronal view, the MNI T1-weighted template (29,30), shown with the locations of waypoint ROIs for classification of the left corticospinal tract (5) (slightly enlarged for visualization purposes). The subject’s anisotropic power map (APM) (31) is used as the target for registration, due to its similarity to the T1 contrast. **C** Classification of the streamlines. Here, in a lateral view, the streamlines classified as belonging to the left corticospinal tract (CST L), overlaid on a mid-saggital slice of the subject’s non diffusion-weighted (b0) image. The streamlines are shaded by the subject’s fractional anisotropy (FA) along their length. **D**, Tract profiles are extracted from the bundles. Here, the FA profile for CST L.

**Table S1.** Abbreviations of the major white matter pathways recognized by pyAFQ.

|  |  |
| --- | --- |
| ARC L | Left Arcuate |
| ARC R | Right Arcuate |
| ATR L | Left Thalamic Radiation |
| ATR R | Right Thalamic Radiation |
| CGC L | Left Cingulum Cingulate |
| CGC R | Right Cingulum Cingulate |
| CST L | Left Corticospinal |
| CST R | Right Corticospinal |
| FA | Callosum Forceps Minor |
| FP | Callosum Forceps Major |
| IFO L | Left Inferior Fronto-occipital Fasciculus |
| IFO R | Right Inferior Fronto-occipital Fasciculus |
| ILF L | Left Inferior Longitudinal Fasciculus |
| ILF R | Right Inferior Longitudinal Fasciculus |
| SLF L | Left Superior Longitudinal Fasciculus |
| SLF R | Right Superior Longitudinal Fasciculus |
| UNC L | Left Uncinate |
| UNC R | Right Uncinate |

**Fig. S2.**
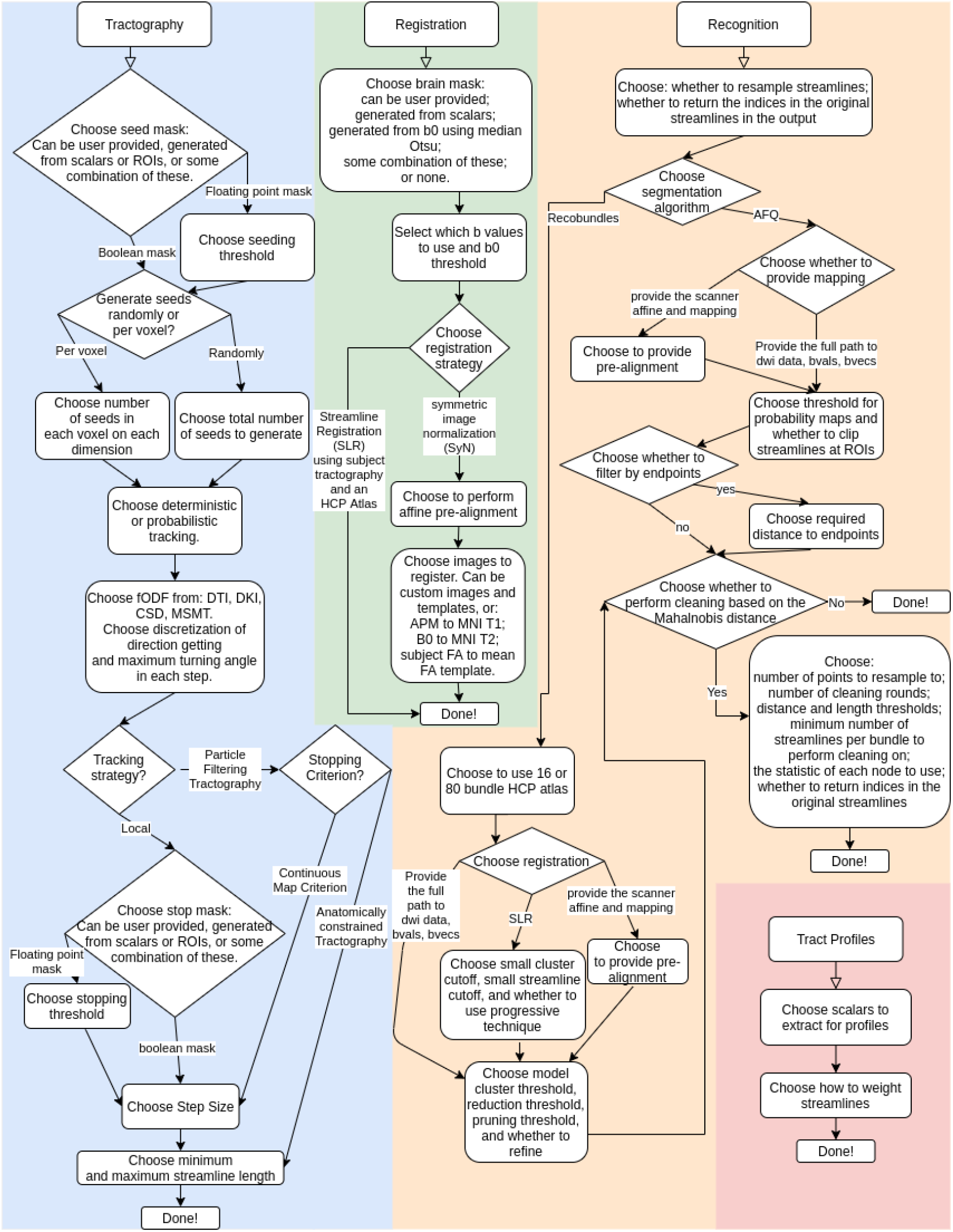
Choices the user can make for how to run pyAFQ. The colors represent different steps of tractometry. Tractography is shaded blue, registration is shaded green, recognition is shaded orange, and tract profiles is shaded red. Every rounded box and diamond contains one or more choices, except for the rounded boxes marked “Done!”, which indicates all choices have been made. Diamonds indicate the path you take depends on the choice in the diamond. pyAFQ has reasonable defaults for all of these decisions; however it also makes it simple for the user to customize their tractometry pipeline according to their needs.

**Fig. S3.**
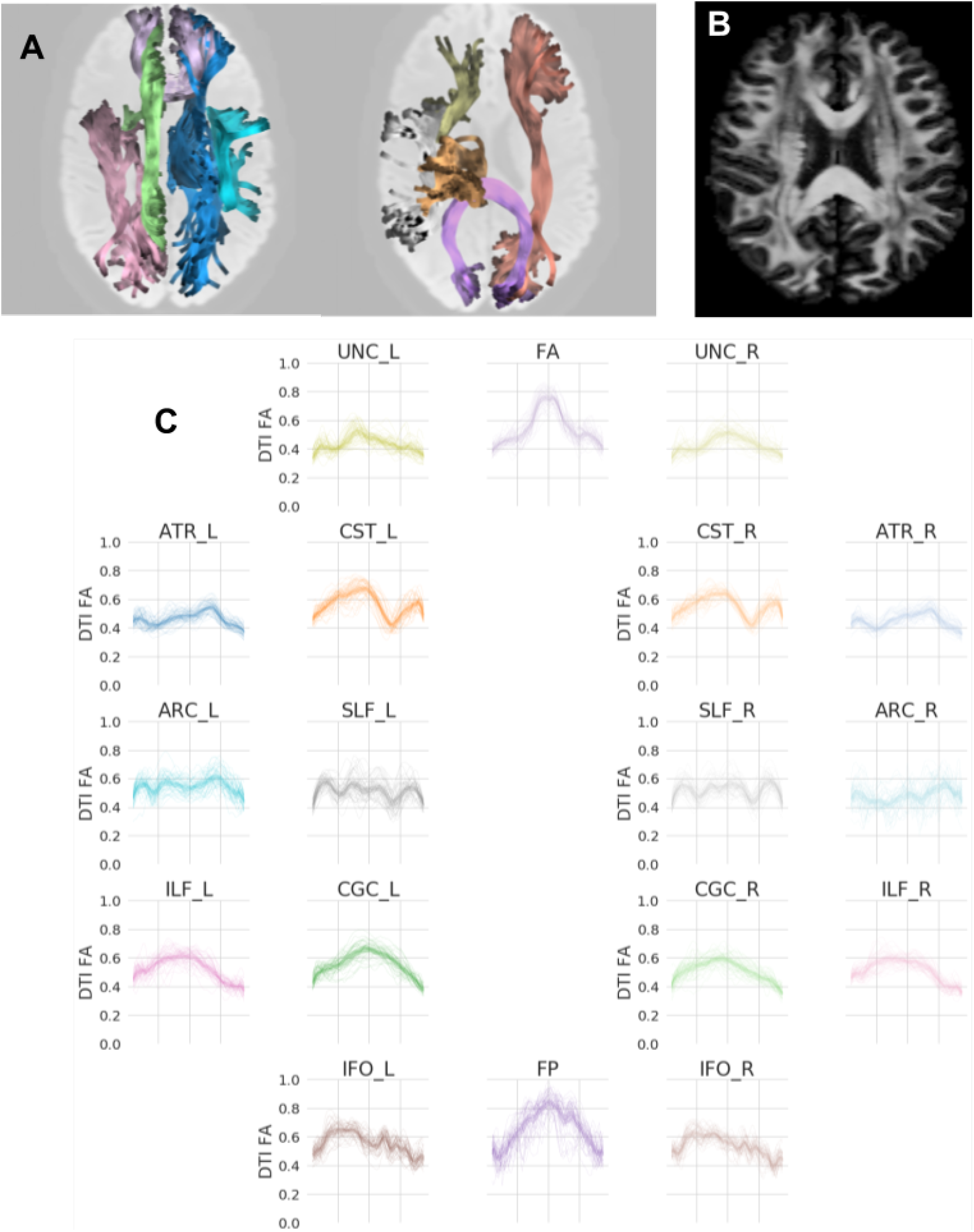
Extraction of tract profiles from the recognition of white matter into major bundles of streamlines. **A** Representative bundles from an example subject in the HCP-TR dataset. Streamlines are colored by bundle, and are shaded by the interpolated FA value at each point. The background is the mean non diffusion-weighted image (b0). **B** The same subject’s fractional anisotropy (FA). **C** extracting FA along each bundle and plotting the FA in a tract profile. Individual tract profiles are plotted with thin lines and the mean tract profile is plotted with a thick line. The tract profiles are colored according to their bundle are laid out in positions that reflect their anatomical positions (compare **A** and **C**).

**Fig. S4.**
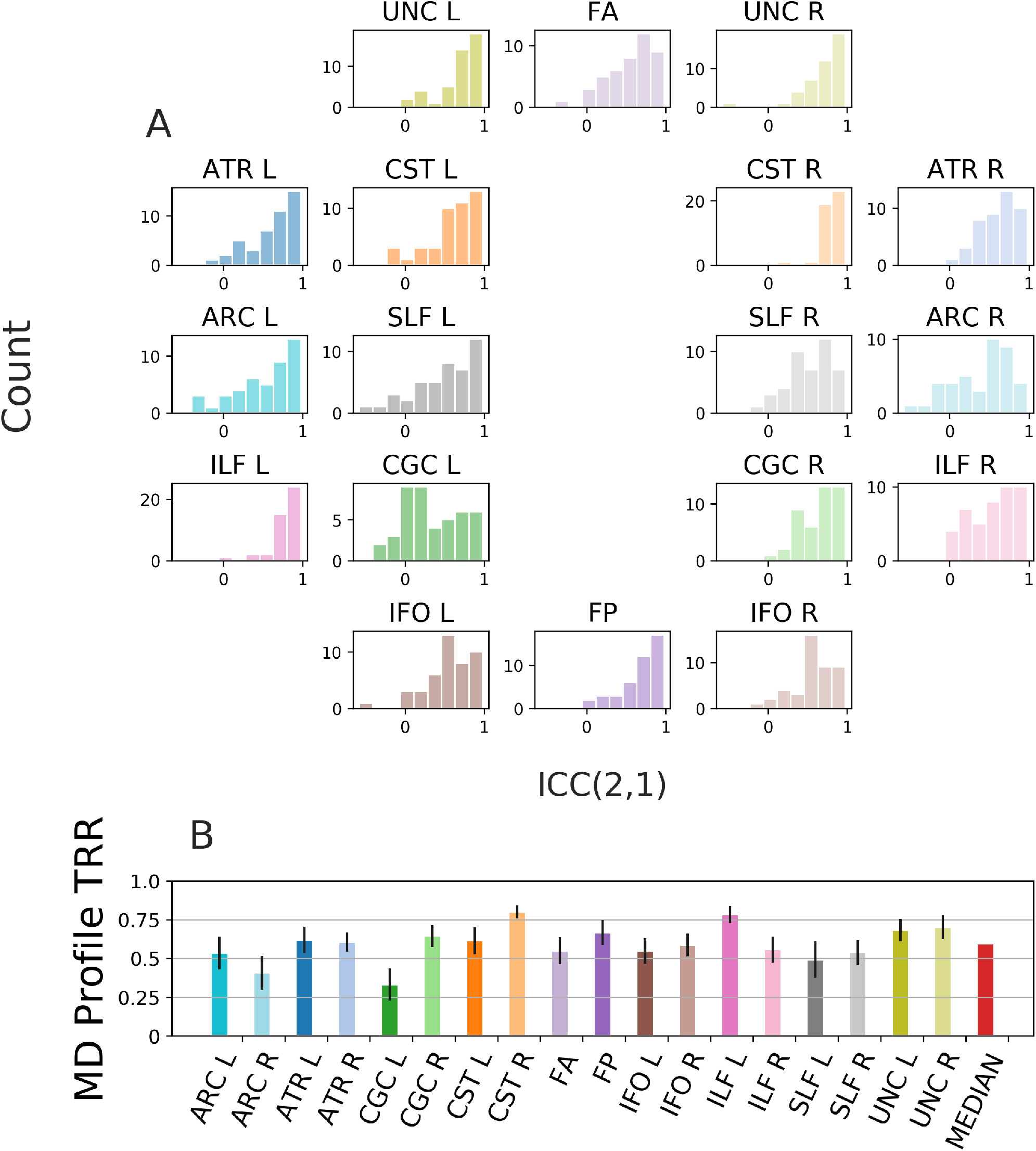
MD profile test-retest reliability. **A:** Histograms of individual subject ICC between the MD tract profiles across sessions for a given bundle. Colors encode the bundles, matching the diagram showing the rough anatomical positions of the bundles for the left side of the brain (center). **B:** Mean (±95% confidence interval) TRR for each bundle, color-coded to match the histograms and the bundles diagram, with median across bundles in red.

**Fig. S5.**
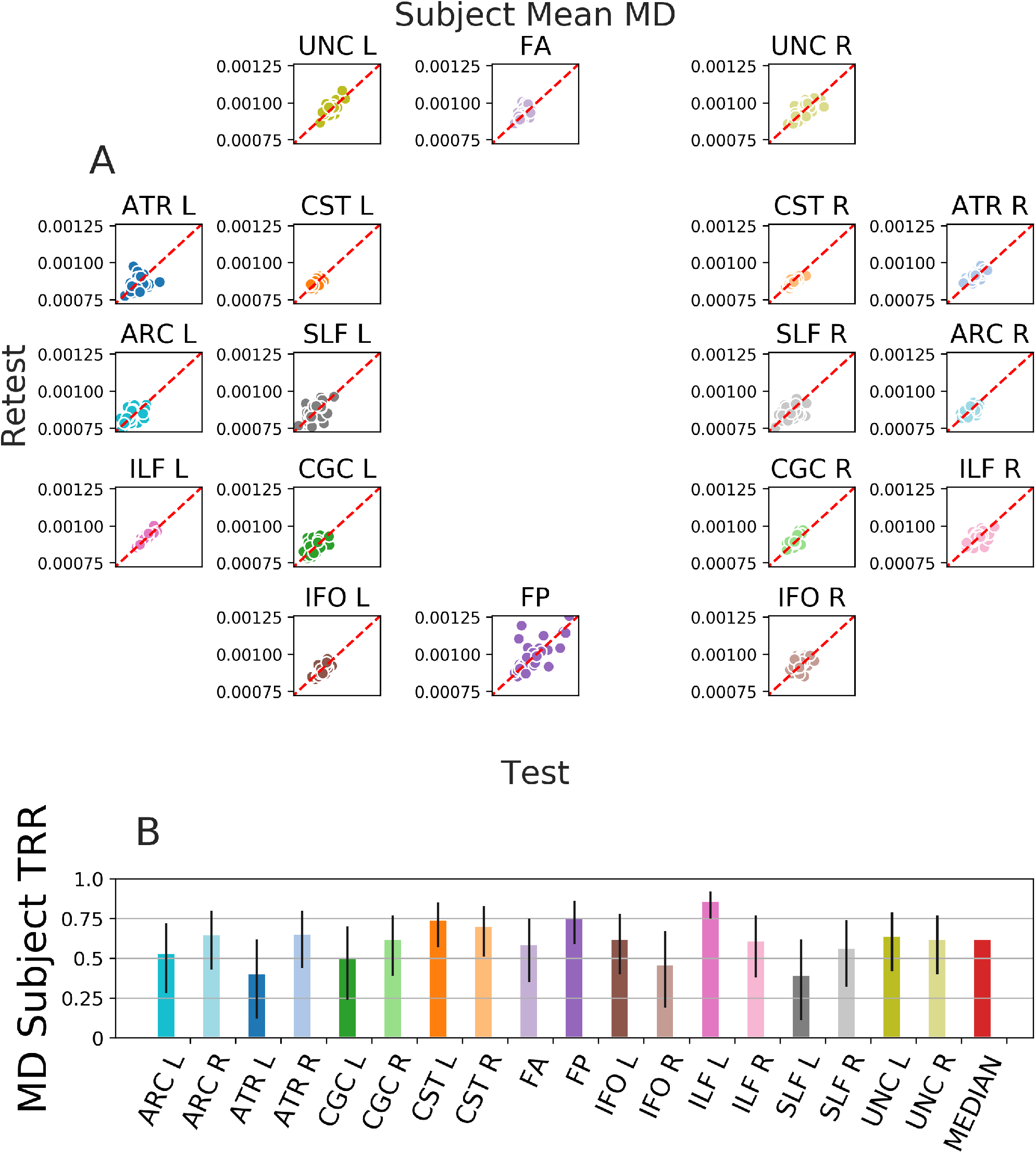
Subject test-retest reliability. **A:** Mean tract profiles for a given bundle and the MD scalar for each subject using the first and second session of HCP-TR. Colors encode bundle information, matching the core of the bundles (center). **B:** subject reliability is calculated from the Spearman’s *ρ* of these distributions, with median across bundles in red. Error bars show the 95% confidence interval.

**Fig. S6.**
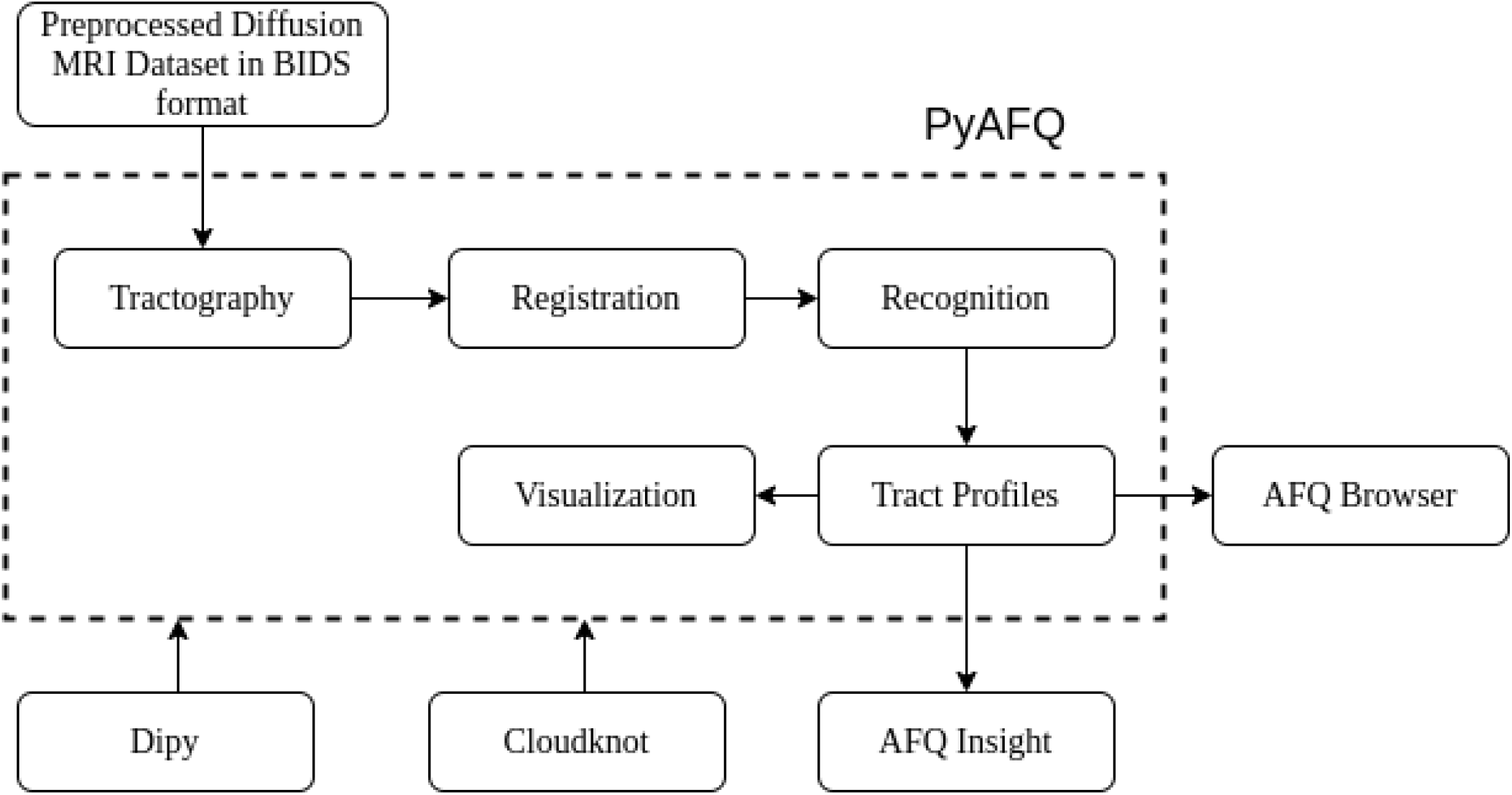
The pyAFQ software is intergrated into an ecosystem for reproducible tractometry. Steps performed by pyAFQ are enclosed in the dotted rectangle, whereas steps outside that rectangle are performed by other software. Upper left: pyAFQ requires preprocessed diffusion MRI data in BIDS format. This could be from QSIprep (26) or dMRIprep (https://github.com/nipreps/dmriprep). Bottom right: pyAFQ outputs can serve as inputs to AFQ Browser for further interaction and visualization (52) or AFQ Insight for statistical analysis (20). Bottom left: pyAFQ uses DIPY (28) for the implementation of dMRI algorithms. pyAFQ uses Cloudknot (63) to scale processing by parallelizing across subjects in AWS.

**Fig. S7.**
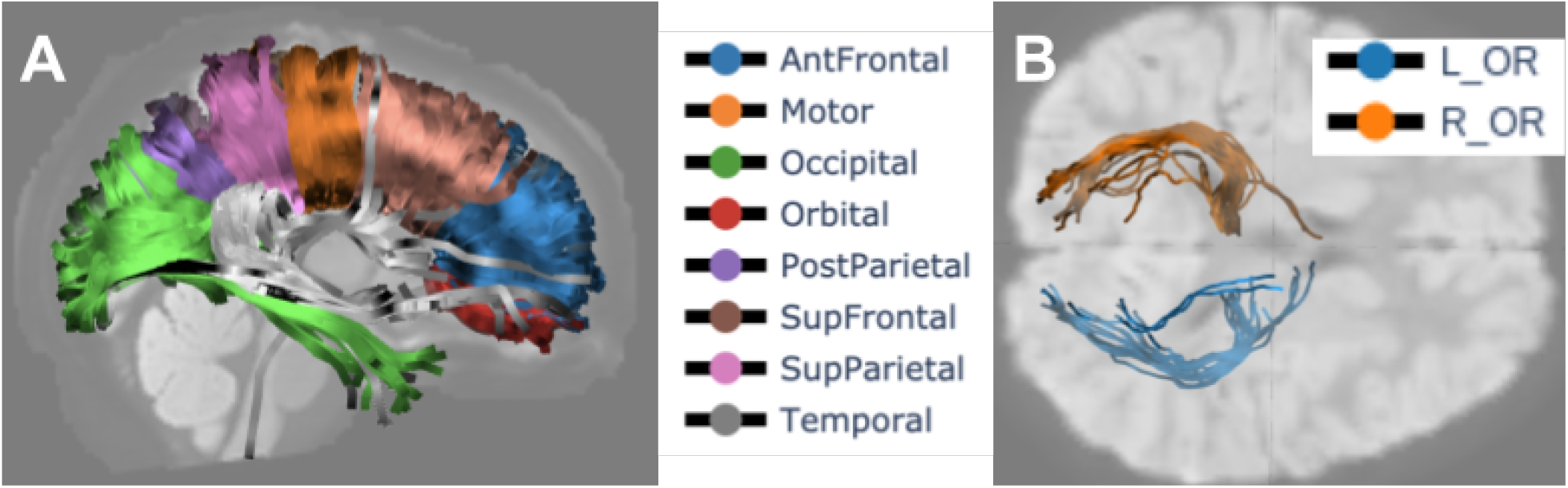
Callosal bundles from HCP-TR, optic radiations from UW-PREK, found by pyAFQ. Streamlines are colored according to their bundles and shaded according to FA. The background images are each a b0 slice. **A** callosal bundles found by pyAFQ on an example subject from HCP-TR. **B** optic radiations found by pyAFQ on an example subject from UW-PREK.

